# KlebSeq: A Diagnostic Tool for Healthcare Surveillance and Antimicrobial Resistance Monitoring of *Klebsiella pneumoniae*

**DOI:** 10.1101/043471

**Authors:** Jolene R. Bowers, Darrin Lemmer, Jason W. Sahl, Talima Pearson, Elizabeth M. Driebe, David M. Engelthaler, Paul Keim

## Abstract

Healthcare-acquired infections (HAIs) kill tens of thousands of people each year and add significantly to healthcare costs. Multidrug resistant and epidemic strains are a large proportion of HAI agents, and multidrug resistant strains of *Klebsiella pneumoniae*, a leading HAI agent, have become an urgent public health crisis. In the healthcare environment, patient colonization of *K. pneumoniae* precedes infection, and transmission via colonization leads to outbreaks. Periodic patient screening for *K. pneumoniae* colonization has cost-effective and life-saving potential. In this study, we describe the design and validation of KlebSeq, a highly informative screening tool that detects *Klebsiella* species and identifies clinically important strains and characteristics using highly multiplexed amplicon sequencing without a live culturing step. We demonstrate the utility of this tool on several complex specimen types including urine, wound swabs and tissue, several types of respiratory, and fecal, showing *K. pneumoniae* species and clonal group identification and antimicrobial resistance and virulence profiling, including capsule typing. Use of this amplicon sequencing tool can be used to screen patients for *K. pneumoniae* carriage to assess risk of infection and outbreak potential, and the expansion of this tool can be used for several other HAI agents or applications.

## INTRODUCTION

*Klebsiella pneumoniae* has been a leading healthcare-acquired infection (HAI) agent for decades (1, 2). Emergence of multidrug-resistant *K. pneumoniae*, especially expanded-spectrum β-lactamase (ESBL) producers and carbapenemase producers, has elevated the morbidity and mortality rates and healthcare costs associated with *K. pneumoniae* to highly significant levels (3-6). Healthcare-and outbreak-associated strain types of *K. pneumoniae* that appear highly transmissible and have a propensity for antimicrobial resistance (AMR) or virulence gene acquisition are a growing proportion of the *K. pneumoniae* species (7-18). ST258, the crux of the worldwide carbapenemase-producing Enterobacteriaceae (CPE) threat, disseminated rapidly around the world’s healthcare systems despite its recent emergence (17). Its progenitor strains in clonal group (CG) 258 also cause outbreaks and carry many important ESBL and carbapenemase genes (9, 19-21). Several other strain types such as those in CG14, CG20, and CG37, also frequently appear as multidrug resistant and in outbreak situations (7, 10, 12, 15).

Host colonization is likely an important reservoir driving transmission of these strains. In the healthcare environment, intestinal colonization of *K. pneumoniae* is risk factor for infection (22-24), and carriers of CPE are at high risk for invasive disease (25). Rates of CPE and ESBL-producing *K. pneumoniae* colonization are rising in patient and healthcare worker populations, increasing the size of the reservoir, and increasing chances of transmission (26, 27). Asymptomatic transmission of multidrug resistant strains is rapid (16, 28), and transmission events that lead to outbreaks often go undetected (29, 30). Early detection of *K. pneumoniae* colonization of healthcare patients, especially multidrug resistant *K. pneumoniae* or epidemic strain type colonization, is now considered critical to infection control (24, 30-33).

Infection control programs that include detection and isolation of carriers have repeatedly been successful in markedly decreasing multidrug resistant or epidemic strain infections (31, 34-37), but this practice is uncommon. Many of these programs use culture-based methods such as antibiotic-containing broth enrichment or selective media for detecting CPE or ESBL producers, which have several limitations including turn-around time, narrow application, lack of sensitivity and specificity, subjectivity, and extensive labor for high-throughput screening (31, 38). Automated systems also require up-front organism culture and isolation, with many of the same limitations (31). PCR-based assays are rapid, but often use DNA from culture, or if used on DNA extracted from specimens, may have low sensitivity. Additionally, a limited number of tests can be run simultaneously, and may miss important AMR genes not previously known to circulate in a given locale (31, 39).

In this study, we describe a new tool, KlebSeq, for screening and surveillance that detects and characterizes *Klebsiella* from complex samples such as wound and nasal swabs or fecal samples without culturing using relatively easy-to-use multiplex amplicon sequencing. KlebSeq includes a sizeable panel of assays for species identification, strain identification, and important virulence and AMR gene targets designed to generate information for hospital epidemiology and infection prevention. Results from screening a patient population with this system would rule in or rule out the possibilities of particular transmission events, and identify patients carrying high-risk strains like ST258 or other multidrug resistant *Klebsiella*. The highly multiplexed nature of the amplicon sequencing tool greatly expands the capacity of a single sequencing run, minimizing costs, and allows for high-throughput patient sample testing. Additionally, this innovation can serve as a model system for many other applications including targeting other HAI agents and their multiple AMR mechanisms.

## METHODS

### Samples

Isolates for target identification and assay validation, along with DNA extracted from clinical specimens were acquired through collaborations with a large hospital reference laboratory that receives specimens from ten system-wide medical centers in Arizona, and from a high volume private reference laboratory that receives specimens from regional inpatient, long-term care, and outpatient facilities. Clinical specimen types included various respiratory specimens (nasal, ear, and throat swabs, sputa, tracheal aspirates, and bronchial alveolar lavages), urine, and wound swabs or tissue. DNA was extracted from isolates by Qiagen DNeasy Blood and Tissue Kit with additional lytic enzymes when appropriate. DNA was extracted from clinical specimens by NucliSENS easyMAG (bioMerieux, Durham, NC). DNA from healthy donor fecal samples was acquired from a family microbiome study. Samples had been collected from members of seven families over multiple timepoints. DNA was extracted following the Earth Microbiome Project protocol (40).

### Assay target identification and assay design

Figure 1 illustrates the methodologies and resources utilized, also described below, to amass a target library and develop several types of amplicon sequencing assays.

**Figure 1.**
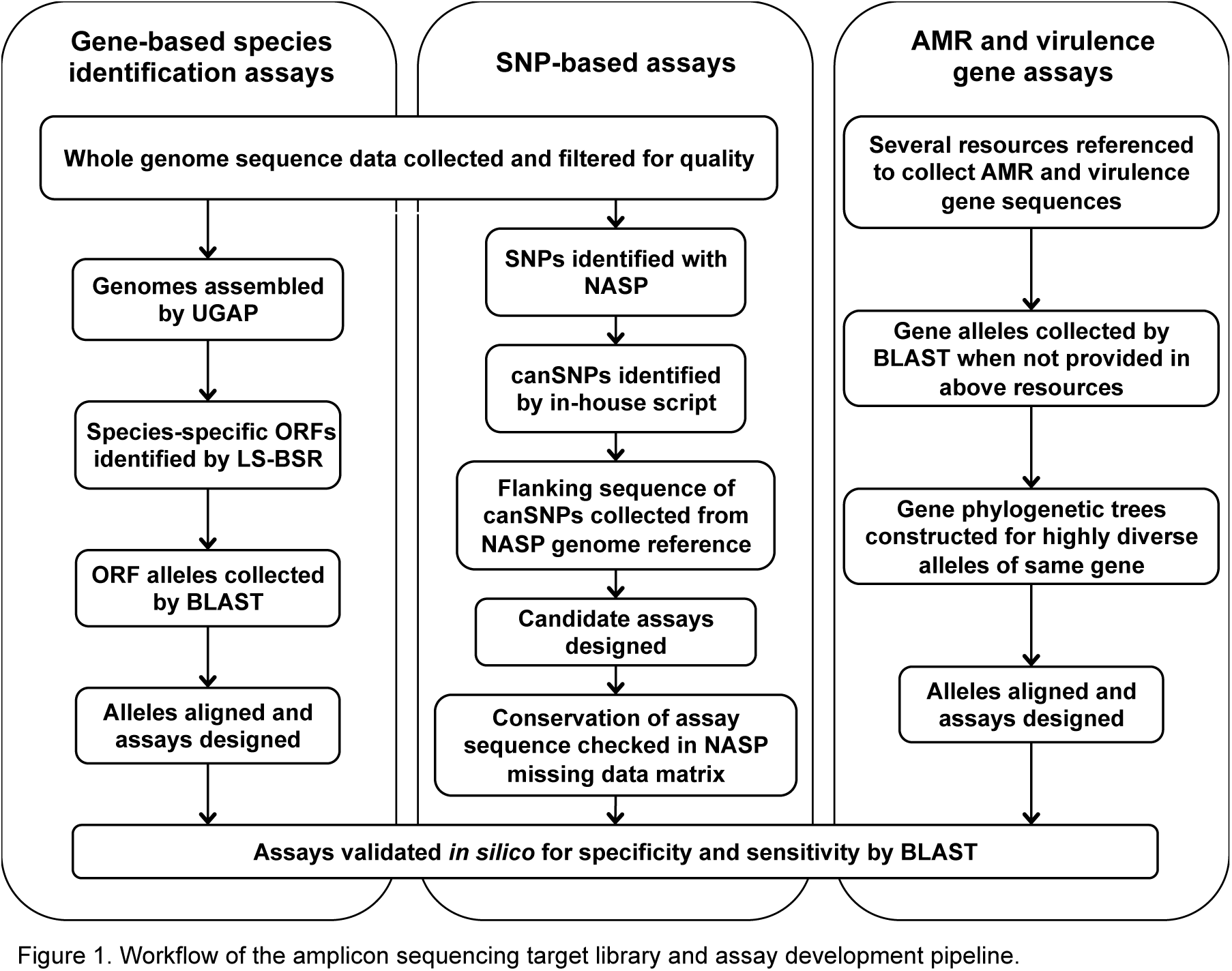
Workflow of the amplicon sequencing target library and assay development pipeline.

### Whole genome sequencing, SNP detection, and phylogenetic analysis

In-house genome libraries were prepared from 130 *Klebsiella* isolates with a 500 base pair insert size using KAPA Library Preparation Kit with Standard PCR Library Amplification (Kapa Biosystems, Wilmington, MA) and sequenced on Illumina’s GAIIx or MiSeq. Public genome sequence data from 277 *K. pneumoniae*, 18 K. *quasipneumoniae*, and 13 *K. variicola* isolates were downloaded from the SRA database (http://www.ncbi.nlm.nih.gov/Traces/sra/) and 178 *K. pneumoniae*, four *K. quasipneumoniae*, and 11 *K. variicola* from the Assembly database (http://www.ncbi.nlm.nih.gov/assembly), and all passed filters for high quality. The publicly available bioinformatic tool NASP (41), developed for microbial genome analysis, was used to detect single nucleotide polymorphisms (SNPs) among the genomes. In brief, reads were aligned to a reference genome, either one from concatenated scgMLST alleles (10) or MGH 78578 (Genbank accession no. CP000647) using Novoalign (Novocraft.com) and SNPs were called with GATK (42). Data filtered out included SNP loci with less than 10X coverage or with less than 90% consensus in any one sample, regions duplicated in the reference genome as identified by Nucmer, and SNP loci that were not present in all genomes in the dataset. In NASP, results were output in a SNP matrix from a core genome common to all isolates in the analysis. Phylogenetic trees were generated from the NASP SNP matrices using MEGA 6.0 (43) and subsequently plotted by means of ITOL v2 or v3 (44).

### Genomic target identification

To find whole gene targets for assay design, select genomes were assembled with UGAP (https://github.com/iasonsahl/UGAP), which uses the SPAdes genome assembler (45). Assemblies were then run through LS-BSR (46), which generates a list of ORFs that have high identity among target species genomes and that have low identity or are not present in non-target genomes. Alleles of the candidate target ORFs were collected by BLAST, including alleles from non-target genomes if present. Lastly, alleles of candidate ORFs were aligned for assay design. Two ORFs (M1 and M2) were collected for two *K. pneumoniae*-specific assays. Canonical SNPs (canSNPs) were identified from the SNP matrix generated from NASP. Sequence flanking each SNP was collected from the NASP reference genome.

### AMR and virulence gene target collection

AMR and virulence gene sequences were identified and collected in several ways, including from http://www.lahey.org/studies/other.asp#table1, http://www.lahey.org/qnrstudies/, the *Klebsiella* BIGSdb http://bigsdb.web.pasteur.fr/klebsiella/, public literature, and NCBI http://www.ncbi.nlm.nih.gov. Public literature included Holt *et al*. (47), in which a species-wide analysis of *K. pneumoniae* genomes revealed several siderophore systems and other virulence factors more associated with infectious than colonizing strains. AMR genes include the major ESBL and carbapenemase genes, plasmid-mediated quinolone resistance determinants as well as the *gyrA* and *parC* chromosomal genes, several aminoglycoside resistance genes, TMP-SMX, tetracycline, streptomycin, chloramphenicol, and fosfomycin resistance genes, and the recently discovered plasmid-mediated colistin resistance gene *mcr-1*. Virulence targets include several siderophore systems, for which multiple genes from each were used as assay targets, the regulator of mucoid phenotype (an indicator of a hypervirulence), and *wzi* gene for capsule typing, for which we used the published assay (48), and two genes highly associated with invasive infection pK2044_00025 and pK2044_00325 (47). For genes that consist of highly diverse alleles, for example *bla*_CTX-M_, *qnrB*, or *dfrA*, phylogenetic trees based on nucleotide sequences were generated in order to group similar alleles for assay design.

### Assay design and validation

Gene-based target alleles were aligned in SeqMan (DNAStar, Madison, WI) to identify conserved regions for primer design, and assays were designed with guidance from RealTimeDesign™ (Biosearch Technologies, Petaluma, CA), or gene-based assays were generated with AlleleID^®^ (Premier Biosoft, Palo Alto, CA), which designs assays to capture alleles in an alignment rather than to individual sequences. SNP assay primers were designed using RealTimeDesign™, and primer sequences were checked for conservation in the NASP SNP matrix. Lastly, assays were run through BLAST http://blast.ncbi.nlm.nih.gov/Blast.cgi, to check for cross-reactivity to other relevant targets or species, including human. Universal tails were added to each primer sequence for library preparation, as described in Amplicon library preparation below. The assays and their primer sequence are listed in Table S1.

Individual assays were screened across positive controls when accessible, and screened across several isolate gDNAs to increase confidence in robustness, especially when known positive controls were not available. Additionally, multiplex PCR was validated by initial gene-specific PCR (described below) followed by PCR product dilution, then screening of individual assays by Sybr green-based qPCR. For this, 10 μL reactions of 1X Platinum SYBR Green qPCR SuperMix (ThermoFisher Scientific, Waltham, MA), 200 nM forward and reverse primers of one assay, and 1 μL diluted multiplex PCR product were run at 95°C initial denaturation for 4 min, then 40 cycles of 95°C for 15 s and 60°C for 1 min. Lastly, several panels of known isolate DNAs were screened with the amplicon sequencing method to test for sensitivity and specificity of the species and strain identification assays. AMR and virulence gene assays were validated by comparing amplicon sequencing results with whole genome sequence data.

### Amplicon library preparation and sequencing

Amplicon library preparation using universal tails was described in detail previously (49). Here, assays were pooled for multiplex PCR. Initial gene-specific PCR comprised 12.5 μL Kapa Multiplex PCR Mastermix (Kapa Biosystems, Wilmington, MA), 10 μL primer mix (for final concentration of 200 nM each), and 2.5 μL DNA template from each sample, and was denatured at 95°C for 3 min, cycled 25 times at 95°C 15 s, 60°C 30 s, 72°C 1 min 30 s, with final extension 72°C 1 min. Multiple multiplex PCR products from the same sample were pooled, and PCR product pools were cleaned with 1X Agencourt AMPure XP beads (Beckman Coulter, Indianapolis, IN). A second PCR using the universal tail-specific primers added Illumina’s sample-specific index and sequencing adapters. This PCR comprised 12.5 μL 2X Kapa HiFi HotStart Ready Mix (Kapa Biosystems), 400 nM each primer, and 1 to 10 μL cleaned gene-specific PCR product, and was denatured 98°C 2 min, cycled 6 to 12 times at 98°C 30 s, 65°C 20s, 72°C 30 s, with final extension 72°C 30 s. Final PCR product was cleaned with 0.8X Agencourt AMPure XP beads (Beckman Coulter).

Amplicon libraries from individual samples were quantified by qPCR using Kapa Library Quantification Kit (Kapa Biosystems). Samples were then pooled in equimolar concentration for sequencing on the Illumina MiSeq platform with 2x250bp version 2 kit.

### Analysis

Amplicon sequencing results were automatically analyzed using a newly developed amplicon sequencing analysis pipeline (ASAP) (50), which uses a JavaScript Object Notation (JSON) file that describes all assays in the multiplex. Information in the JSON file included a category for each assay (presence/absence, SNP, gene variant, or region of interest) that dictates how ASAP will report results, and reference sequences for mapping. In ASAP, amplicon sequence reads were first trimmed of adapter or read-through sequences with Trimmomatic (51), then mapped to the reference sequences with an aligner. BAM alignment files were analyzed alongside the JSON file assay descriptions to determine presence, breadth and depth of coverage, and proportions of nucleotide polymorphisms for each amplicon. User defined parameters included use of the bowtie2 aligner (52), and thresholds for determining results of screening included 80% breadth and 100X depth of coverage for isolate DNA, and 80% breadth and 20X depth of coverage and ≥10% proportion of polymorphism for informative SNP loci for complex specimen DNA (meaning that at least 10% of the reads had to share a SNP state at a given locus for it to be reported). ASAP output included an XML file containing details of the analysis of each assay target for each sample, which can be converted into a webpage interface using XSLT transformations. SeqMan NGEN (DNAStar, Madison, WI) and Tablet (53) were used to verify results.

## RESULTS

### Phylogenetic analysis and canSNP identification

Using the *Klebsiella* strict core genome MLST (scgMLST) (10) assembly as a reference, SNPs among a diverse set of genomes from *K. pneumoniae* and genomes from the newly defined *K. quasipneumoniae* (22 from the public databases and one from in-house isolates) and *K. variicola* (24 from the public databases and five from in-house isolates) were identified with NASP. A canonical SNP that differentiates *K. quasipneumoniae* and one that differentiates *K. variicola* from *K. pneumoniae* were selected for assay development.

Using the reference genome MGH 78578 and 548 diverse *K. pneumoniae* genomes, NASP generated a SNP matrix from which canonical SNPs for each of the major clonal groups were selected for assay development. Clonal groups and locations of canSNPs identifying 35 clonal groups and sequence types in the context of the *K. pneumoniae* species are illustrated in Figure 2. Redundancy was intentionally included in the assays expected to be positive for the most epidemic strains of *K. pneumoniae* such as ST14, ST20, and strains in CG258 in order to increase confidence in positive results.

**Figure 2.**
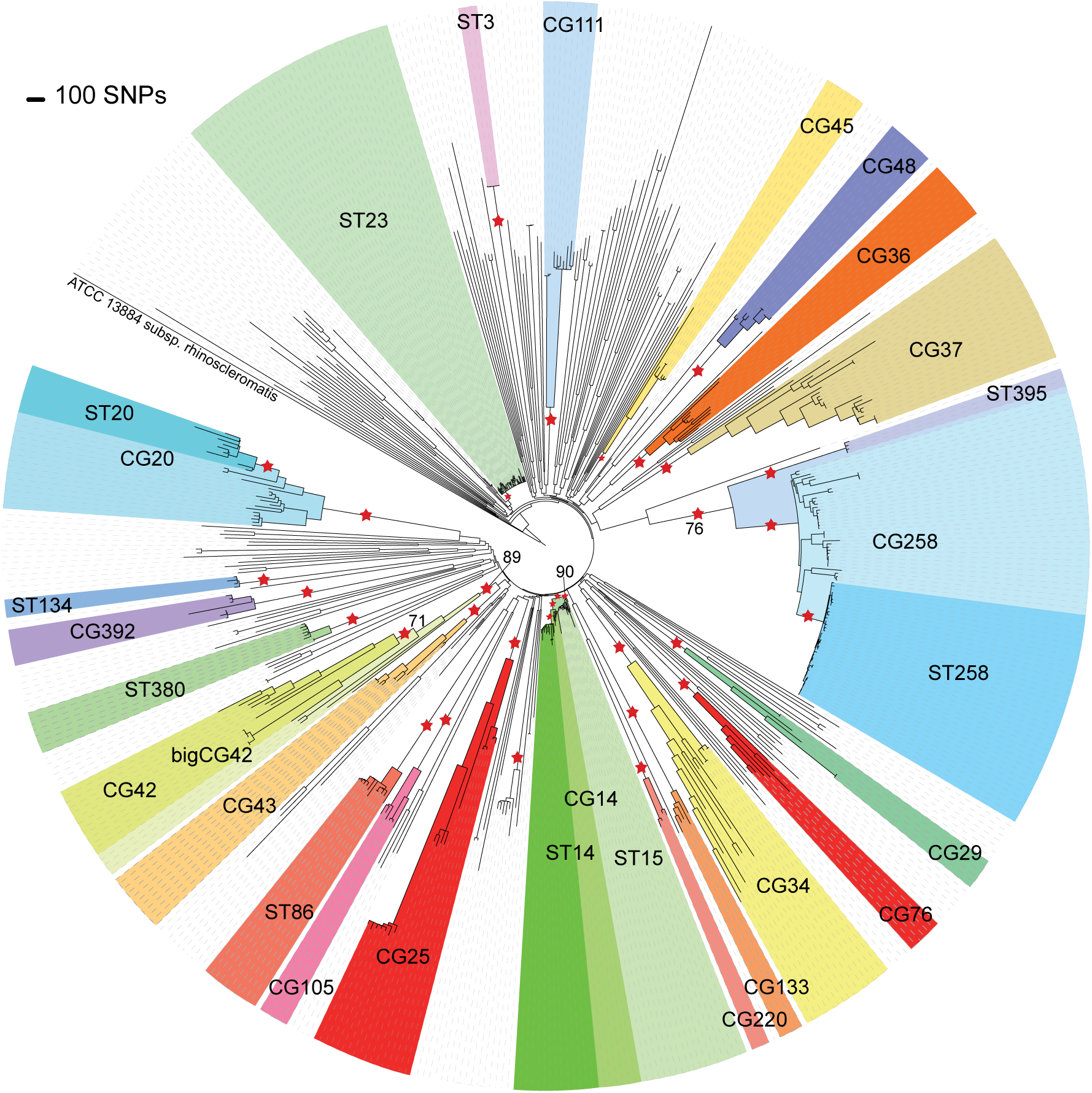
Maximum parsimony tree with 100 bootstraps of the SNPs among 548 *K. pneumoniae* genomes. Major clonal groups are colored, and locations of canonical SNPs for strain identification assays are marked with stars. All branches labeled with canonical SNPs had >99% bootstrap support, except on the three branches indicated.

### KlebSeq validation

For validation of the species and clonal group identification assays, genomic DNA from 73 *K. pneumoniae* isolates that had also been whole genome sequenced (4 of which were later identified as *K. quasipneumoniae* and *K. variicola*, see below), 22 *K. oxytoca* isolates, and 157 other enteric, opportunistic-pathogen isolates, which included *E. coli, Enterobacter aerogenes, E. amnigenus, E. cloacae, E. hormaechei, Enterococcus faecalis, E. faecium, E. sp*., *Proteus mirabilis, Providencia stuartii*, and *Serratia marcescens* were screened with KlebSeq. Sensitivity and specificity results of the species identification assays compared with clinical microbiological identification are in Table 1. With the redundancy built into the multiplex by including two assays Kp-M1 and Kp-M2, which target two different *K. pneumoniae* species-specific genes (M1 and M2), 100% sensitivity is achieved. One isolate previously identified as *K. pneumoniae* typed as *K. quasipneumoniae* and two as *K. variicola*. These isolates’ whole genomes were added to the phylogenetic analysis of these three species that was previously run to find the species-specific canSNPs (see Methods). The *K. quasipneumoniae* and *K. variicola* genomes identified by our assay clustered with their respective species in the phylogeny (Fig. 3). Clinical methods do not currently distinguish among all three of these species, so assay sensitivity and specificity were not calculated for *K. quasipneumoniae* and *K. variicola*.

**Table 1.** Sensitivity and specificity of the KlebSeq species identification assays on genomic DNA from known isolates, DNA from urine, wound, and respiratory specimens for which clinical culture results are known, and results of screening across specimens of unknown content.

| Isolate DNA<br>n=252 | Total<br>screened | Amplicon sequencing assay<sup>a</sup> |  |  |  |  |  |
| --- | --- | --- | --- | --- | --- | --- | --- |
|  |  | Kp-M1 | Kp-M2 | Kp-M1 + Kp-M2 | Kquasi | Kvari | Koxy |
| *K. pneumoniae* | 69 | 68 | 67 | 69 | 0 | 0 | 0 |
| *K. quasipneumoniae* | 2 | 0 | 0 | 0 | 2 | 0 | 0 |
| *K. variicola* | 2 | 2 | 0 | 0 | 0 | 2 | 0 |
| *K. oxytoca* | 22 | 0 | 0 | 0 | 0 | 0 | 21 |
| Non-target species | 157 | 2/88 | 0/88 | 2/88 | 0/155 | 0/155 | 0/135 |
| Sensitivity |  | 99% | 97% | 100% | 100% | 100% | 95% |
| Specificity |  | 98% | 100% | 98% | 100% | 100% | 100% |
| Urine DNA<br>n=46 | Total<br>screened | Kp-M1 | Kp-M2 | Kp-M1 + Kp-M2 | Kquasi | Kvari | Koxy |
| *K. pneumoniae* | 16 | 14 | 16 | 16 | 2 (1 mix<sup>b</sup>) | 1 (mix<sup>b</sup>) | 0 |
| *K. oxytoca* | 6 | 1 | 1 | 1 (CG34) | 0 | 0 | 6 |
| Other species | 24 | 2 | 2 | 2 (1 CG392) | 0 | 0 | 0 |
| Unknown | 0 | 0 | 0 | 0 | 0 | 0 | 0 |
| Sensitivity |  | 88% | 100% | 100% | - | - | 100% |
| Specificity |  | 90% | 90% | 90% | - | - | 100% |
| Wound DNA<br>n=40 | Total<br>screened | Kp-M1 | Kp-M2 | Kp-M1 + Kp-M2 | Kquasi | Kvari | Koxy |
| *K. pneumoniae* | 1 | 0 | 0 | 0 | 0 | 1 | 0 |
| *K. oxytoca* | 1 | 0 | 0 | 0 | 0 | 0 | 1 |
| Other species | 31 | 3 | 3 | 3 (1 CG29) | 0 | 0 | 0 |
| Unknown | 7 | 1 | 1 | 1 | 0 | 0 | 1 |
| Sensitivity |  | 100% | 0% | 100% | - | - | 100% |
| Specificity |  | 94% | 94% | 94% | - | - | 100% |
| Respiratory specimen DNA<br>n=87 | Total<br>screened | Kp-M1 | Kp-M2 | Kp-M1 + Kp-M2 | Kquasi | Kvari | Koxy |
| *K. pneumoniae* | 6 | 6 | 6 | 6 | 0 | 0 | 0 |
| *K. oxytoca* | 1 | 0 | 0 | 0 | 0 | 0 | 1 |
| Other species | 77 | 7 (1<br>ST258) | 5 | 7 (2 CG37, 1<br>CG36) | 0 | 0 | 1 |
| Unknown | 3 | 0 | 0 | 0 | 0 | 0 | 0 |
| Sensitivity |  | 100% | 100% | 100% | - | - | 100% |
| Specificity |  | 92% | 95% | 92% | - | - | 99% |
| Fecal specimen DNA<br>n=89 | Total<br>screened | Kp-M1 | Kp-M2 | Kp-M1 + Kp-M2 | Kquasi | Kvari | Koxy |
| Unknown | 89 | 8 | 5 | 8 | 1 (mix<sup>b</sup>) | 3 (2 mix<sup>b</sup>) | 13 |

<sup>a</sup> Assay descriptions are as follows: Kp-M1 and Kp-M2 are *K. pneumoniae* species identification assays that detect targets M1 and M2 in the *K. pneumoniae* genome. Kquasi, Kvari, and Koxy are the *K. quasipneumoniae*-, *K. variicola*-, and *K. oxytoca*-specific assays, respectively.
<sup>b</sup> These species were found as mixtures with *K. pneumoniae*, based on a proportion ( $\geq 10\%$ ) of sequencing reads containing the species-defining SNP.

**Figure 3.**
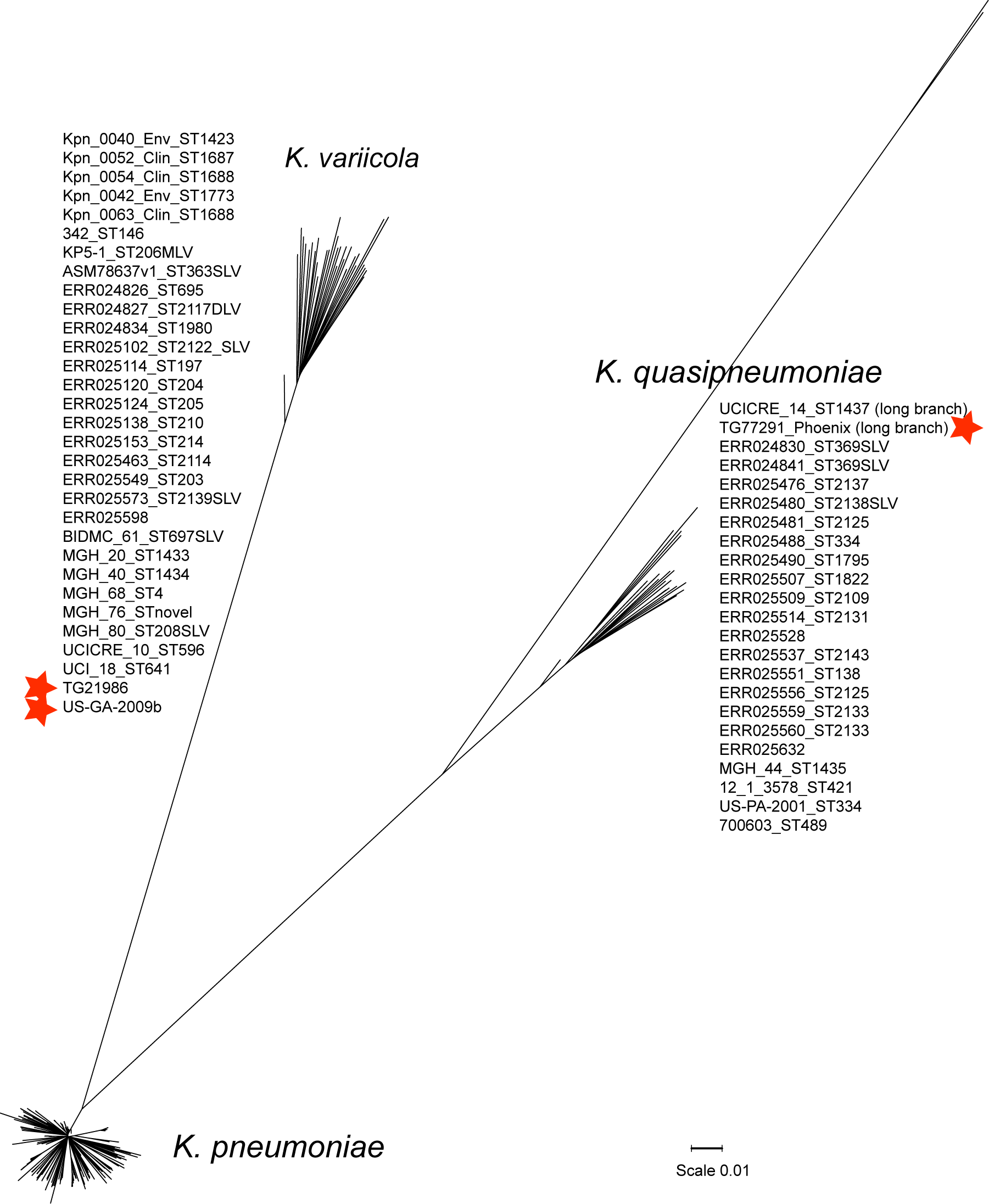
Neighbor-joining tree with 100 bootstraps of the SNPs among the diverse set of *K. pneumoniae*, the *K. variicola*, and *K. quasipneumoniae* genomes, with our unknown specimens that typed as *K. variicola* and *K. quasipneumoniae* labeled with stars.

Table 2 shows results of the *K. pneumoniae* clonal group identification and capsule typing assays. Each isolate’s strain type was correctly captured by the appropriate assays, or not captured in cases where no assay was designed for that clonal group.

**Table 2.** Isolates used for assay validation and results of strain typing with amplicon sequencing.

| Isolate sequence type | No. isolates | ASAP Strain Typing Assay Results | Capsule Typing Results by Partial *wzi* Sequence<sup>a</sup> |
| --- | --- | --- | --- |
| ST11 | 3 | CG258, CG258wo395 | wzi-39 or -75, wzi-74, not typable |
| ST14 | 5 | CG14, ST14, innerST14 | wzi-2 |
| ST14 SLV | 1 | CG14 | wzi-16 |
| ST15 | 2 | CG14, ST15 | wzi-24 or -45 |
| ST20 | 2 | CG20, ST20 | wzi-84, wzi-118 |
| ST23 | 8 | ST23 | wzi-1 |
| ST34, ST34 SLV | 2 | CG34 | wzi-114, wzi-12 |
| ST36 | 2 | CG36 | wzi-27 or -79 |
| ST37 | 2 | CG37 | wzi-50, wzi-39 or -75 |
| ST39 | 1 | No group | wzi-2 |
| ST42 | 2 | bigCG42, CG42 | wzi-29 |
| ST43 | 1 | CG43 | wzi-30 |
| ST45 | 1 | CG45 | wzi-133 |
| ST65 | 1 | CG25 | wzi-72 |
| ST101 | 2 | CG43 | wzi-29, wzi-137 |
| ST107 | 1 | No group | wzi-74 |
| ST111 | 1 | CG111 | wzi-63 |
| ST147 | 1 | CG392 | wzi-64 |
| ST152 | 1 | CG105 | wzi-150 |
| ST228 | 1 | CG34 | wzi-116<sup>b</sup> |
| ST234 | 1 | No group | wzi-114 |
| ST249 | 2 | No group | wzi-128 |
| ST258 group, no clade | 6 | CG258, CG258wo395, 3xST258 | wzi-154 |
| ST258 group, clade 2 | 2 | CG258, CG258wo395, 3xST258, clade 2 | wzi-154 |
| ST277 | 1 | No group | wzi-97 or -185 |
| ST334 | 1 | *K. quasipneumoniae* | wzi-68 |
| ST340 | 2 | CG258, CG258wo395, ST340 | wzi-50, wzi-173 |
| ST376 | 1 | bigCG42, CG42 | wzi-2 |
| ST380 | 1 | ST380 | wzi-203 |
| ST636 | 1 | No group | wzi-155 |
| ST719 | 1 | No group | wzi-192 |
| ST776 | 1 | No group | wzi-39<sup>b</sup> or -75<sup>b</sup> or -193<sup>b</sup> |
| ST833 | 1 | CG258, CG258wo395 | wzi-50 |
| ST978 | 1 | *K. quasipneumoniae* | wzi-212<sup>b</sup> |
| ST1401 | 1 | No group | wzi-96 |
| ST82 | 2 | No group | wzi-128 |
| ST260 | 1 | No group | wzi-1 |
| ST360 SLV | 1 | *K. variicola* | wzi-53 |
| ST427 SLV | 1 | No group | wzi-64 |
| ST513 SLV | 1 | No group | wzi-87 |
| ST815 SLV | 1 | No group | wzi-114<sup>b</sup> |
| ST244 SLV | 1 | No group | wzi-162<sup>b</sup> |
| ST2006 | 1 | *K. variicola* | wzi-227 |
| ST2055 | 1 | No group | wzi-14 |

<sup>a</sup> The Illumina version 2 chemistry used provides approximately 500 bp of sequence data. The amplicon size for the *wzi* assay (48) is approximately 580 bp.
<sup>b</sup> *wzi* allele represents best match when one or multiple SNPs were present.

Partial sequencing of the *wzi* gene for capsule typing gave surprisingly clear results, given that approximately 75 bp of the informative region is missing in our sequence output, as it was based on a previously published assay (48). The amplicon size of the *wzi* PCR is approximately 580 bp, which is too long to cover with the Illumina version 2 sequencing chemistry. However, our data show the promise of full capsule typing by *wzi* sequence with longer read chemistry *(i.e*. Illumina version 3 chemistry, which provides 2x300 bp reads). Results from screening non-target organisms showed that several of the *K. pneumoniae* clonal group assays amplified DNA from other organisms, as was expected, however all SNP states that define a particular clonal group are specific to that clonal group. As such, sequence analysis by ASAP reported when a clonal group was present only if the defining canSNP state was present, and reported nothing if it was not.

AMR gene detection by amplicon sequencing was validated by comparing ASAP results with AMR gene screening of whole genome sequence with SRST2 (54). Results showed good correlation with few discrepancies, which were almost all negative by amplicon sequencing while positive by SRST2. This may have been due to PCR primer discrepancy or inefficiency or failure in amplicon library preparation. Only six samples account for a majority of the negatives, which points to the latter alternative. Virulence gene detection was validated by comparing ASAP results from whole genome sequence data with those from amplicon sequence data. Results showed almost perfect concordance. In addition, by targeting genes that are part of the same virulence factors (i.e. siderophore systems), sensitivity and confidence in results was increased.

These results also confirm that KlebSeq is applicable to pure isolates as well as complex specimens. Screening isolate DNA has the added benefit of traceability of the AMR and virulence genes, which are often carried on mobile genetic elements, to their host. Isolate screening could be used for surveillance and other purposes for identifying or characterizing *Klebsiella*.

### Specimen sample results

KlebSeq was run on DNA from 87 respiratory specimens, 46 urine specimens, 40 wound specimens, and 89 fecal samples from healthy individuals. Sensitivity and specificity results of the species identification assays compared with clinical microbiological methods are shown in Table 1. In most cases, sensitivity was very high, except in the wound specimens where the one sample clinically identified as *K. pneumoniae* typed as *K. variicola*. Amplicon sequencing identified several samples with *Klebsiella* that went undetected with clinical microbiological methods, including several in which clonal groups were also detected.

Important clonal groups of *Klebsiella* were detected in multiple specimens. In the 18 urine samples positive for *K. pneumoniae*, clonal identifications included CG34, ST20, CG45, CG392 (which includes the NDM-producer ST147 (55), though these samples were negative for *bla*_NDM_, n=2), ST133, and CG111. In wounds, one strain identification, CG29, was made from the three *K. pneumoniae*-positive samples. From respiratory specimens, groups CG37 (n=2), ST134 (n=1), ST258 (n=2), CG36 (n=3), and innerST14 (n=1) were identified. Interestingly, several clonal groups were identified in the healthy donor fecal specimens as well. Out of the eight *K. pneumoniae-positive* samples, groups included ST20 (an alignment of which is shown in Figure 4), CG37, and CG76, which are all members of multidrug-resistant outbreak strain types (11, 12, 15), along with ST133 and ST380. ST380 is associated with a K2 capsule type and hypervirulence, and causes pyogenic liver abscesses in healthy people, especially of Asian ethnicity (56). Many Asians are colonized with hypervirulent, K1 or K2 capsule strain types; however the level of risk of subsequent liver infection is unknown (56). For this sample, no *wzi* gene sequence was obtained, thus the capsule type is unknown. A majority of the *K. pneumoniae* in our samples did not fall into the major clonal groups targeted by KlebSeq. Likely these strains all belong to lesser-known clonal groups, as more studies are showing that many *K. pneumoniae* infections are caused by non-epidemic, sporadic strains (57, 58).

Numerous and variable AMR genes were detected in the specimens, including different variants of the same gene that confer different phenotypes. Using sequence-based information, we demonstrate that many of the *K. pneumoniae* had key mutations in the *gyr*A gene known to confer resistance to fluoroquinolones. Additionally, several samples contained the *aac(6’)-Ib* gene for aminoglycoside resistance, and some of those contained the minor sequence variant *aac(6’)-Ib-cr* for fluoroquinolone resistance; mixtures of these two genes were also detected. A majority of the infection specimens (non-healthy donor specimens), both positive and negative for *K. pneumoniae*, were positive for other aminoglycoside resistance genes, as well as tetracycline, TMP-SMX, streptomycin, fosfomycin, and chloramphenicol resistance genes. A few contained plasmid-mediated quinolone resistance genes. Several samples, especially the respiratory specimens, were also positive for KPC and CTX-M groups 1 and 9 genes. Most of the healthy donor specimens were positive for TMP-SMX resistance genes, and many for streptomycin, aminoglycoside, tetracycline, and fosfomycin resistance genes. Some also contained plasmid-mediated quinolone resistance genes. Fortunately, none were found to contain ESBL or carbapenemase genes. No complex specimens in the study were positive for genes encoding the important carbapanemases OXA-48, VIM, or NDM, and none were positive for the plasmid-mediated colistin resistance gene *mcr-1*.

These sets of samples did not appear to contain especially virulent strains of *K. pneumoniae*. The yersiniabactin siderophore genes were by far the most prevalent of the virulence genes tested, although positive samples made up less than half of the *K. pneumoniae*-positive samples (46%). No samples were positive for *rmpA*, regulator of mucoid phenotype gene, including the ST380-containing sample, and few were positive for the salmochelin siderophore genes, which are associated with invasive *K. pneumoniae* infection (47). One respiratory specimen that contained a ST14 strain was positive for a K2 capsule type by partial *wzi* sequencing. K2 strains of *K. pneumoniae* are associated with hypermucoviscosity and hypervirulence, as previously mentioned. However this respiratory sample was not positive for *rmpA*, and a recent study proposed that the presence of multiple siderophore system genes (linked to K1 or K2 capsule genes) explains hypervirulence rather than capsule type (47). In our data, *K. pneumoniae*-containing samples were positive for multiple siderophores or other virulence-associated genes only 15% of the time. Sequencing of *wzi* revealed a variety of capsule types, and incidences where the same clonal groups had different *wzi* genotypes and where they had the same genotype. This character would help identify or rule out a transmission event when patients carrying the same strain are found.

On an interesting note, in the healthy donor fecal samples collected from members of the same families over time, out of the eight *K. pneumoniae*-positive samples, only two came from the same person over time. The characterization assays suggest that the same strain of *K. pneumoniae* was present at both time points. The fact that there were not more cases of positive results from the same person in multiple rather than single time points is interesting. This could be due to intermittent shedding of *K. pneumoniae* in feces, intermittent colonization of *K. pneumoniae*, or heterogeneity in the sample itself, underrepresenting the full microbial community when a small sample is taken. *K. pneumoniae*-positive samples were found in multiple members of two of the seven families. In one of these families, the positive members carried different strains from one another, and in the other it appears two members shared a CG37 with the same capsule type. The sample set is too small to draw conclusions from the data; however, the data raise interesting questions about community *K. pneumoniae* carriage.

## DISCUSSION

The frequency of HAI in the United States is estimated at one in 25 hospital patients, totaling hundreds of thousands, with significant mortality (2). HAIs have a significant impact on healthcare costs; a 2009 CDC report estimated upwards of $45 billion in annual additional cost (59). Infections of AMR organisms cause significantly higher mortality rates and ICU admissions, and significant excess healthcare costs including hospitalization, medical care, and antimicrobial therapy over infections of susceptible strains (5, 60). HAI prevention measures, although costly in and of themselves (61), have the potential to save many lives and billions of dollars (59). Periodic patient screening and isolation of AMR organism carriers have proven successful in controlling transmission and outbreaks in several hospitals (31, 34-37). Use of a highly informative screening and surveillance tool such as KlebSeq has cost-effective and life-saving potential.

Early detection of *K. pneumoniae* colonization of healthcare patients, especially multidrug resistant *K. pneumoniae*, would allow healthcare staff to make more informed patient management decisions. In outbreak situations, rapid identification of transmissions before subsequent infections would allow for proactive measures to curb an outbreak. In non-outbreak situations, identification of particular strains and AMR genes would help to assess the risk of *K. pneumoniae* carriage to the host patient as well as to other patients, as some strains are more associated with outbreaks, HAI, AMR, and treatment failure than others (7-13, 62). Likewise, identification of virulence genes also informs risk, as particular virulence factors are more associated with pathogenic than colonizing *K. pneumoniae* (47). Additionally, many *K. pneumoniae* infections, including HAIs and non-multidrug resistant infections, are caused by non-epidemic, lesser-known strain types (57, 58). Classifying the *K. pneumoniae* in each patient sample would help an institution decide when and which intervention procedures should be enacted, and also understand more about transmission dynamics and local strain type circulation.

The amplicon sequencing assay and analysis pipeline described here has several characteristics that make it ideal as a healthcare screening tool. With a single assay, enough information is garnered about a patient’s *Klebsiella* carriage status to contribute greatly to patient management or to infection control decisions. Indexing samples by means of the universal tail during sample preparation allows characterization of a large number of specimens in one run, minimizing sequencing costs per specimen and allowing for high-throughput screening of hundreds of patient samples simultaneously. KlebSeq uses DNA extracted directly from a specimen so targets from entire populations of a species are analyzed to capture different strains in the same sample, which can be numerous (63, 64). If culture-based methods are used for screening, different strains are missed when one genotype *(i.e*. colony) is chosen for characterization, and resulting information is limited. Additionally, culture-based methods can miss “silent” multidrug resistant *K. pneumoniae* that test negative for carbapanemases *in vitro* (16), and if used for high-throughput screening, they can be laborious, time-consuming, costly, and subjective (31, 38). If screening of large numbers of patients by amplicon sequencing is cost-prohibitive, it can be limited to the highest-risk groups of patients, *i.e*. long-term care facility patients (31, 65), travelers returning from endemic regions (66, 67), ICU patients (28), patients that previously carried (67-69), patients that shared a room with a known carrier (70) or case contacts of carriers (71), those who’ve recently taken antibiotics (72, 73), or patients on mechanical ventilation, enteral feeds, or that have had prior *Clostridium difficile* infections (74). Additionally, using ASAP makes the analysis in KlebSeq streamlined, and results are easily interpretable. Lastly, the amplicon sequencing and ASAP surveillance approach is customizable and updateable. Individual assays can be added or removed, adding only the cost of new primers.

The results we present here show that KlebSeq works on DNA from numerous sample types, including pure organism culture, complex, multi-organism samples, and swab samples with low-level microbial DNA in a presumably high human DNA background. Our data show that in addition to identifying different species of *Klebsiella*, clinically important clonal lineages of *K. pneumoniae* can be identified from culture or complex specimens without culture methods. Our method distinguishes *K. pneumoniae* from *K. quasipneumoniae* and *K. variicola*, the latter of which appear to be lower risk species with regard to infection and virulence (47), and we identify cases where these species were previously misidentified as *K. pneumoniae*. We highlight several instances where culture methods failed to produce a positive *K. pneumoniae* result, including one sample that contained the critical ST258 strain. We identified dozens of AMR and virulence genes within individual samples, demonstrating the additional function of profiling for clinically important characteristics, and were able to distinguish minor genotype differences that confer different phenotypes, i.e. the *gyrA* gene, *aac(6’)-Ib* versus *aac(6’)-Ib-cr*, and the *wzi* gene.

Notably, our data show that several healthy individuals carry clinically important strains of *K. pneumoniae* as well as many AMR genes and siderophore virulence systems. For our purposes, these healthy donor fecal DNA samples were used to validate the usage of our amplicon sequencing approach on highly complex gut metagenome samples. Much more study is needed to elucidate the implications of healthy host carriage of known pathogenic strains of *K. pneumoniae*. Furthermore, the fact that we observed carriage of the hypervirulence-associated ST380 strain from a healthy person, and the hypervirulence-associated K2 capsule type in a ST14 strain from a respiratory infection, lends credence to the idea that we need much more information about avirulent *K. pneumoniae* to be able to draw conclusions about these associations.

Overall, the KlebSeq method was able to accurately and consistently identify and characterize *Klebsiella* from complex specimens. A limitation to our study is that clonal group identification in the complex specimens was not confirmed by either whole metagenomic sequencing or from isolation and whole genome sequencing of the *Klebsiella* from the specimen. Additionally, profiling complex specimens directly for AMR and virulence genes, most of which are on mobile elements, can be confounding, as it can’t be known which organism carries the genes. However, KlebSeq is designed for screening and surveillance for high-risk situations using a rule-in/rule-out determination of the possibility of transmission events and through identification of high-risk multidrug resistant or epidemic strains of *Klebsiella*. For these purposes, KlebSeq is ideal. The specimen types used to validate could be considered a limitation, as we did not test rectal swabs, a common specimen type for CPE surveillance, due to unavailability. However, we show KlebSeq works on different swab types and fecal specimens, which addresses the challenges of detection in rectal swabs. Turnaround time from sample collection to result, is dependent only on current technology (not organism culture). We recently ran a proof of concept of a 24-hour sample-to-answer test using different targets (data not shown). This test was done on an Illumina MiSeq using only 60 cycles. Other platforms may allow for this turn around time to be decreased even further Rapid amplicon sequencing with automated analysis and reporting is a promising response to the proposal for constant surveillance for highly transmissible or highly drug resistant pathogens. Our model system, directed at *Klebsiella*, can easily be adapted for multiple other pathogens and for different purposes such as environmental sampling, community host screening, and, as smaller, more on-demand next-generation systems become available, for diagnostics and individual patient monitoring. For several reasons, amplicon sequencing is an applicable tool for healthcare facility surveillance. As these technologies are adopted, considerable coordination within the healthcare facility is paramount to the success of infection and outbreak prevention, with the integration of isolation and barrier precautions, excellent communication, and good stewardship.

Nevertheless, several institutions have shown that the combination of surveillance and systematic response reduces outbreaks and multidrug resistant infections (31, 33-37).

## ACKNOWLEDGEMENTS

We would like to thank Dr. Michael Saubolle of Laboratory Sciences of Arizona for providing isolates and specimen DNA, Krystal Sheridan for technical assistance, and Tricia O’Reilly for administrative support.

